# A nasal vaccine candidate, based on S2 and N proteins from SARS-CoV-2, generates a broad antibody response systemic and at lower respiratory tract

**DOI:** 10.1101/2025.05.09.653001

**Authors:** Yadira Lobaina, Rong Chen, Dania Vazquez-Blomquist, Edith Suzarte, Miaohong Zhang, Zhiqiang Zhou, Yaqin Lan, G Guillen, Wen Li, Yasser Perera, Lisset Hermida

## Abstract

Since the beginning of the COVID-19 pandemic various groups around the world have intensively worked in the development of vaccine candidates against SARS-CoV-2. Several vaccines have been approved in the past years, the majority are based on the Spike or RBD proteins and employs parenteral administration routes. Considering the recent history of Coronavirus zoonotic events, causing serious human health problems, the generation of vaccines with a broad scope of protection and the potential to cut / reduce the transmission remains in the spotlight. The current global pandemic preparedness initiatives have promoted also the preclinical evaluation of a new group of Coronavirus vaccines. In the present work a nasal vaccine candidate based on two highly conserved Sarbecovirus proteins, S2 and nucleocapsid (N), is evaluated in two different mice strains. The vaccine preparation, containing a CpG ODN as adjuvant, was able to generate high antibody titers against both antigens, in sera and bronchoalveolar lavages. This antibody response results cross-reactive to S2 from SARS-CoV-1 and MERS-CoV, and to N from SARS-CoV-1. However, a very low neutralizing capacity was found in the sera of the immunized mice when a pseudoviral system assay was used. On the other hand, the vaccine preparation induces, at systemic compartment, IFN γ secretion, and a marked IgG2a response, specific against both proteins; a profile consistent with the development of a Th1 pattern. Although further evaluations should be done, including protection assays, the demonstrated cross-reactivity level and mucosal response constitutes promising features of this vaccine candidate.

## 1. Introduction

A few years have passed since COVID-19 pandemic onset, a fact that severely impacted the public health worldwide with unprecedented effects. The pandemic posed a big challenge for biomedical research and industry, which obtained, on record time, different vaccines to face the situation. The majority of the COVID-19 approved vaccines are based on the SARS-CoV-2 Ancestral strain Spike protein, or its RBD subunit, with the exception of few inactivated vaccines [**1**]. The Spike protein, and more specifically its S1 subunit containing the RBD, is the main target for the induction of neutralizing antibody response [**2, 3**], which has been correlated with protection [**4, 5**]. However, S1 subunit also constitutes a highly variable region [**6**], then the resulting vaccines show low cross-protective capacity against new SARS-CoV-2 variants or other related Coronavirus [**6**].

More recently, some updating of the original vaccines have been introduced, based on Spike or RBD proteins from the last SARS-CoV-2 variants of concerns like Omicron and other derivatives strain. Lately, the FDA has green-lighted three updated COVID-19 vaccines, Pfizer-BioNTech, Moderna and Novavax [**7**]. Nevertheless, considering the continue SARS-CoV-2 evolution and the real threats of other zoonotic events by Coronavirus that could happen in the future, vaccine candidates with a broader scope of protection are under development [**8**]. There are two main trends in this strategy: to use cocktails preparations, including RBD or Spike proteins from different virus [**9, 10**]; and to include other antigens as vaccine targets, like nucleocapsid and S2 proteins [**11**], considering their high conservation between Coronavirus. Specifically, targeting the S2 subunit constitutes a promising alternative. Some previous works have described the generation of cross-protection using different vaccine candidates based on S2 [**12-18**], the majority of them using parenteral administration and non protein subunit based antigens.

On the other hand, another disadvantage of the former COVID-19 vaccines is their poor capacity to induce immune response at mucosal tissues, like nasal passage and lungs. It is recognized that the presence of specific immune response, antibodies and cell-mediated immunity, at the portal of entry and at the site of infection is crucial to reduce transmission and severity, respectively [**19, 20**]. In the present work, the immune response generated by a vaccine preparation including the full length S2 and N proteins from SARS-CoV-2, and administered by intranasal route, is evaluated using two different mice strains. The cross-reactive profile of the antibody response, generated in sera and bronchoalveolar lavages, against other Betacoronavirus was also studied. So far, no S2-based nasal vaccines are approved, but promising preclinical works highlight their potential for universal coronavirus protection.

## 2. Materials and Methods

### 2.1. Biological reagents

The recombinant antigens were purchased from Sinobiological Inc (China). Nucleocapsid proteins from: SARS-CoV-2, Delta (40588-V07E29) and Omicron (40588-V07E34) variants, and from SARS-CoV-1 (40143-V08B). The S2 protein from SARS-CoV-2 Ancestral strain (40590-V08H1), SARS-CoV-1 (40150-V08B3), and MERS-CoV (40070-V08B).

Specifically, the nucleocapsid protein from Delta strain consisted in amino acids Met1-Ala419 and was expressed in *E. coli*. The S2 protein from Ancestral strain consisted of the extracellular domain, amino acids Ser708-Glu1207, and was expressed in HEK293 cells.

The peptide N_351-365_ from SARS-CoV-2 (ILLNKHIDAYKTFPP) was synthesized with ≥ 97% purity by Zhejiang Peptides Biotech (China).

The ODN-39M, a 39 mer, whole phosphodiester backbone oligodeoxynucleotide (5’-ATC GAC TCT CGA GCG TTC TCG GGG GAC GAT CGT CGG GGG-3’), was synthesized by Sangon Biotech (China).

### 2.2. Vaccine preparation and Immunization experiments

The N protein from SARS-CoV-2 Delta strain was incubated with the ODN-39M, as previously described [**21, 22**]. Briefly, in a 100 µL of reaction, 40 µg of N protein were mixed with 60 µg of ODN-39M in buffer 10 mM Tris, 6 mM EDTA, pH 6.9. The resulting preparations were incubated for 30 min at 30°C in a water bath and then stored at 4°C for 4h. After the preparations were centrifuge at 14000g for 10 minutes. The supernatants were pooled and the protein concentration in the sample was tested. Five micrograms from this preparation were mixed with 10μg of S2 protein from Ancestral strain to obtain the final vaccine preparation dose to be administered per mouse.

Beijing Vital River Laboratory Animal Technology Co., Ltd, and Hunan Prima Drug Research Center Co., Ltd, conducted the mice experiments. Both animal facilities comply with the national standard of the people’s Republic of China GB14925-2010. The immunization protocols were approved by Institutional Animal Care and Use Committee. Six to eight weeks old, females, C57BL/6J (inbred, H-2b) or Balb/C (inbred, H-2d) mice were employed.

All immunogens were dissolved in sterile PBS. For intranasal (in) and subcutaneous (sc) administrations, a volume of 50 μL and 100 μL was employed, respectively. Some immunogens for sc administration contain aluminum hydroxide (alum; Alhydrogel, Invitrogen), as adjuvant, at a final concentration of 1.4 mg/mL.

In a first experiment, C57BL/6J mice were distributed in seven groups of five or more animals each and immunized with three doses, administered on days 0, 7 and 21. The intranasal and subcutaneous routes were evaluated for vaccine administration. Sera samples were collected 18 days after the last dose, and bronchoalveolar lavages (BAL) and spleens were collected 26 days after the last dose. As Placebo controls, a group receiving alum by sc route (PBS+Al sc) and other receiving PBS intranasally (PBS in) were included.

A second experiment was done using Balb/C mice. Groups of six animals were intranasally immunized with three doses, administered on days 0, 15 and 30. Thirty days after the last dose samples of sera, BAL and spleens were collected.

### 2.3. Evaluation of humoral immune response by ELISA

The systemic and mucosal antibody response was evaluated by ELISA. Anti-IgG, subclasses and, -IgA ELISAs were conducted as previously described [**22**]. Briefly, 3 µg/mL of recombinant protein was used to coat 96 well high binding plates (Costar, USA). Plates were subsequently blocked with 2% skim milk solution. Samples were added in duplicates, starting from 1:100 dilution in the case of sera; whereas BALF were directly assayed. Specific horseradish peroxidase conjugates (Sigma, USA) were used. As substrate, OPD (Sigma, USA)/hydrogen peroxide solution was employed. Plates were incubated during 10 min in the dark and the reaction was stopped with 2 N Sulphuric acid. The O.D was read at 492nm in a multiplate reader (FilterMax F3, Molecular Devices, USA). For the evaluations in sera samples the data was represented, mainly, as log10 titers. The arbitrary units of titers were calculated by plotting the O.D values obtained for each sample in a standard curve (hyper-immune serum of known titer). The positivity cut-off was established as 2 times the average of O.D obtained for a pre-immune sera pool. Otherwise, and also for the antibodies detected in BALFs, data were represented as O.D at 492 nm.

### 2.4. IFN-γ ELISPOT

IFN-γ ELISPOT assay was carried out using a specific antibody pair developed for the detection of this cytokine in mice samples (Mabtech, Sweden). Spleen cells were isolated in RPMI culture medium (Gibco, US). Samples (five mice per group) were processed individualized, with the exception of the placebo group for which a pool of three randomly selected mice was evaluated. Duplicates cultures (5x10^5^ and 1x10^5^ splenocytes per well) were incubated for 48 h at 37°C, and 5% CO_2_, in a 96 well round bottom plate (Costar, USA) with a final concentration of 10 µg/mL of each stimulating agent: N351-365 peptide, N or S2 proteins, Concanavalin A (ConA) (Sigma, USA), or medium. The content of the plate was then transferred to an ELISPOT pre-coated plate, and incubated for 16-20 h at 37°C and 5% CO_2_. The successive steps were carried out following the manufacturers recommendations. Finally, the spots were counted using a stereoscopic microscope (AmScope SM-1TSZ, USA) coupled to a digital camera.

### 2.5. Pseudotyped VSV-based neutralization assay

SARS-CoV-2 S protein pseudotyped vesicular stomatitis virus (VSV) based assay was employed to quantify the neutralizing capacity of the sera [**23**]. A commercial kit containing a viral stock of VSV pseudotyped with the whole S protein from Ancestral strain, and a Luciferase substrate solution was acquired (Darui Biotech, China). In a 96 well culture plate (Costar, USA) sera dilutions were incubated with the recommended concentration of virus for 1 h at 37 °C and 5% CO_2_. Two columns of the plates were reserved for virus control (VC, without sera sample), and cells control (CC, without virus). Later, 2x10^4^ Huh-7 cells (cell line provided by Darui Biotech, China) were added to each well and incubated for 24 h at 37°C and 5% CO_2_. After this period, 150 µL of supernatant from each well was removed and 100 µL of Luciferase substrate solution was added. The plates were incubated by 2 min in the dark, and then, the content of each well was resuspended and transferred to a white opaque 96 well plate (Costar, USA). The luminescence was read using a FilterMax F3 microplate reader (Molecular Device, USA). The calculation of the inhibition percentage was done as follows:

Inhibition rate= (1-(average of luminescence for sample – average luminescence for CC)/ (average luminescence for VC – average luminescence for CC)) *100%

Positive and negative sera controls were included in the assay. The positive control consists of a pool of mice sera with a known neutralizing titer 1: 4000. The negative control comprises a pool from the corresponding placebo group. The assay fulfilled with the quality criteria recommended for this kind of test.

### 2.6. Statistical analysis

Graph Pad Prism version 5.00 software (Graph-Pad Software, San Diego, CA, USA) was employed for the statistical analysis. The antibody titers values were transformed to log10 to adjust the data to a normal distribution. For the non sero-converting sera, an arbitrary titer of 1:50 was considered. One-way Anova test was used as parametric tests for multiple group comparisons, followed by a Tukey’s post-test. In the particular cases that required it, non-parametric multiple comparisons using Kruskal Wallis test and Dunns post-tests were employed. The statistical criteria followed the standard considerations for P values: ns, p>0.05; *, p<0.05; **, p<0.01; ***, p<0.001.

## 3. Results

### 3.1 Immunogenicity of the bivalent formulation N+ODN-39M+S2 in C57BL/6J mice

#### 3.1.1 Antibody response in systemic circulation

Eighteen days after the third dose the anti-N and anti-S2 specific IgG and IgG subclasses were measured in sera. As shown in Figure 1, all the animals receiving the bivalent formulation, irrespective of the administration route employed, are able to generate a high IgG response against both proteins.

**Figure 1:**
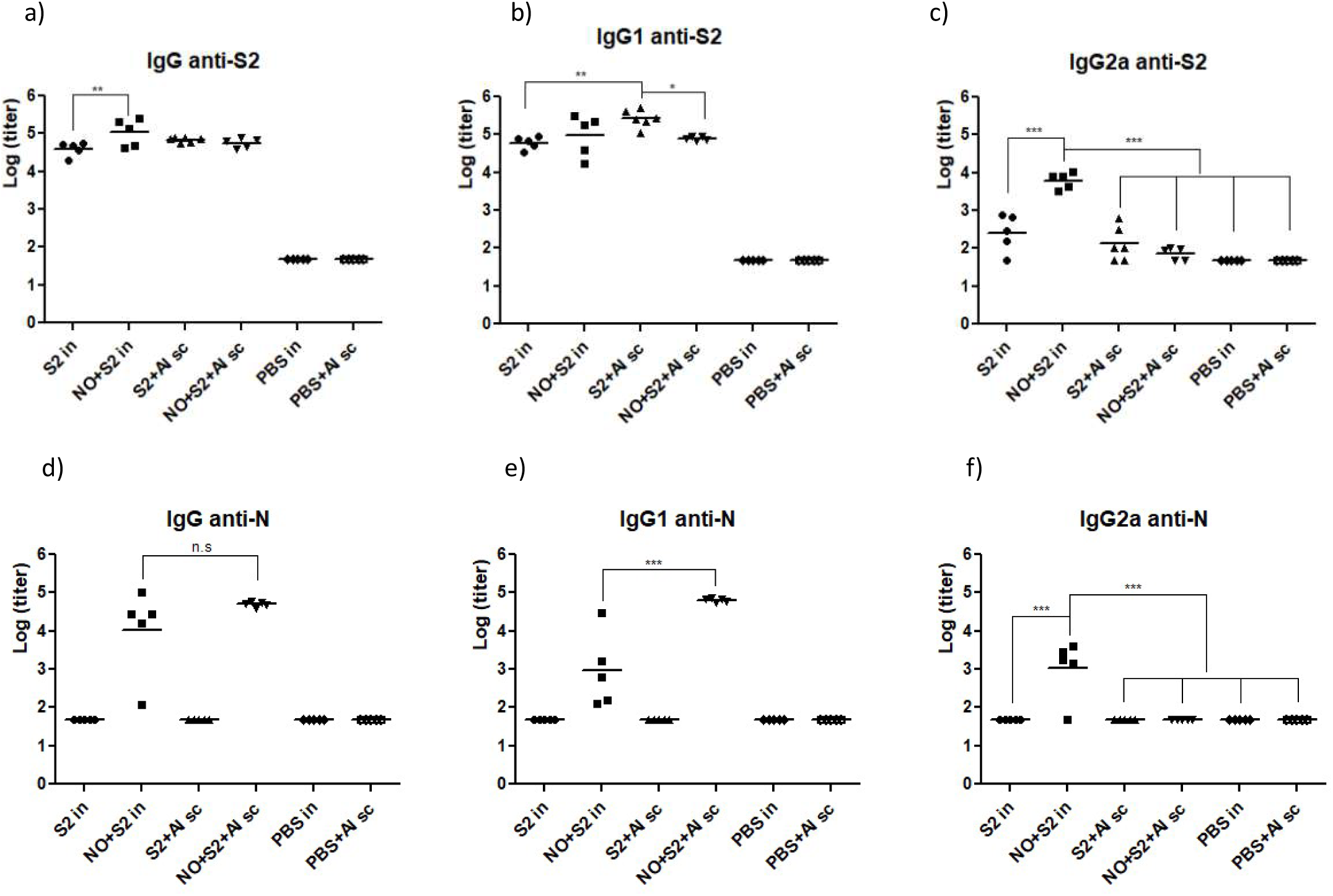
Total IgG and subclasses antibody response measured in sera by ELISA. C57BL/6J mice were immunized with three doses administered at days 0, 7 and 21. A dose of 10μg of S2 protein and 5μg of N protein per mouse was employed. Sera were collected eighteen days after the third dose. Anti-S2 total IgG (a), IgG1 (b) and IgG2a (c). Anti-N total IgG (d), IgG1 (e), and IgG2a (f). NO: N protein + ODN-39M adjuvant, Al: alum, in: intranasal administration, sc: subcutaneous administration.

Interestingly, the S2 protein without adjuvants resulted highly immunogenic by intranasal route, developing high IgG titers after three doses (Figure 1a). However, the inclusion of the mix N+ODN-39M in the preparation is able to significantly increase the anti-S2 titers generated after intranasal administration (p<0.01). On the other hand, the anti-S2 IgG titers induced by intranasal route were similar (p˃0.05) to those induced by subcutaneous administration using preparations that includes alum as adjuvant. Analyzing the S2-specific subclasses response (Figure 1 b and c) it is evident that the increase of IgG observed by intranasal route for the bivalent formulation is associated with the induction of a higher IgG2a titers.

In the case of the anti-N IgG titers (Figure 1d) no differences were observed between the response generated by both groups receiving the bivalent formulation by in or sc routes. However, a higher dispersion in the titers values was evident for the intranasal immunized group. Concerning the anti-N subclasses, the group receiving the bivalent formulation by intranasal route was the only able to induce an IgG2a positive response (Figure 1f).

#### 3.1.2 Specific IgA antibody response in the lower respiratory tract

A positive response of IgA anti-S2 was detected in 50% of the animals that received S2 protein by intranasal route, alone or mixed with N+ODN-39M (Figure 2a). By the contrary the group immunized subcutaneously with the bivalent formulation plus Alum did not show any positive response. Following a similar pattern of response, the IgA anti-N was only detected in the groups immunized by intranasal route, with a trend to develop a higher response for the group receiving the bivalent formulation (Figure 2b).

**Figure 2.**
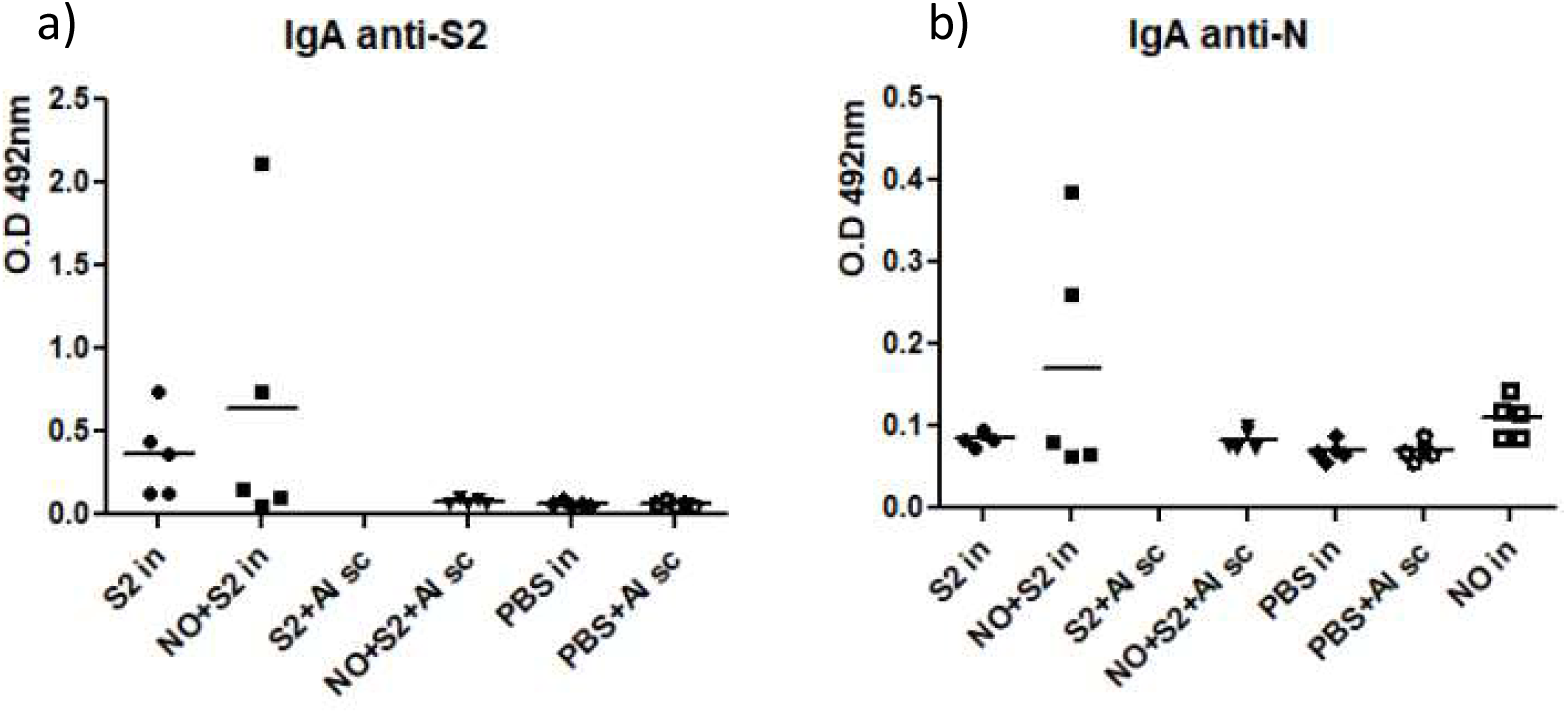
IgA antibody response measured, by ELISA, in bronchoalveolar lavages collected 26 days after the third dose. Anti-S2 (a) and Anti-N (b). C57BL/6J mice were immunized with three doses administered at days 0, 7 and 21. A dose of 10 μg of S2 protein and 5μg of N protein per mouse was employed. No statistical differences were observed among the treatment groups.

#### 3.1.3 Neutralizing capacity of the sera generated after immunization with the bivalent formulation

The bivalent formulation administered by intranasal route induce sera with a low neutralizing capacity (Figure 3a). Four out 5 mice show a positive response, with % of inhibition ≥ 50%, at 1:150 dilution. In the case of the bivalent formulation administered by subcutaneous route with Alum (Figure 3b), the neutralizing capacity induce in the sera was also very low, with 2 out 4 mice with % of inhibition ≥ 50%, at the lower dilution assayed of 1/50; one of these animals showed an inhibition >50% til 1/150 dilution.

**Figure 3.**
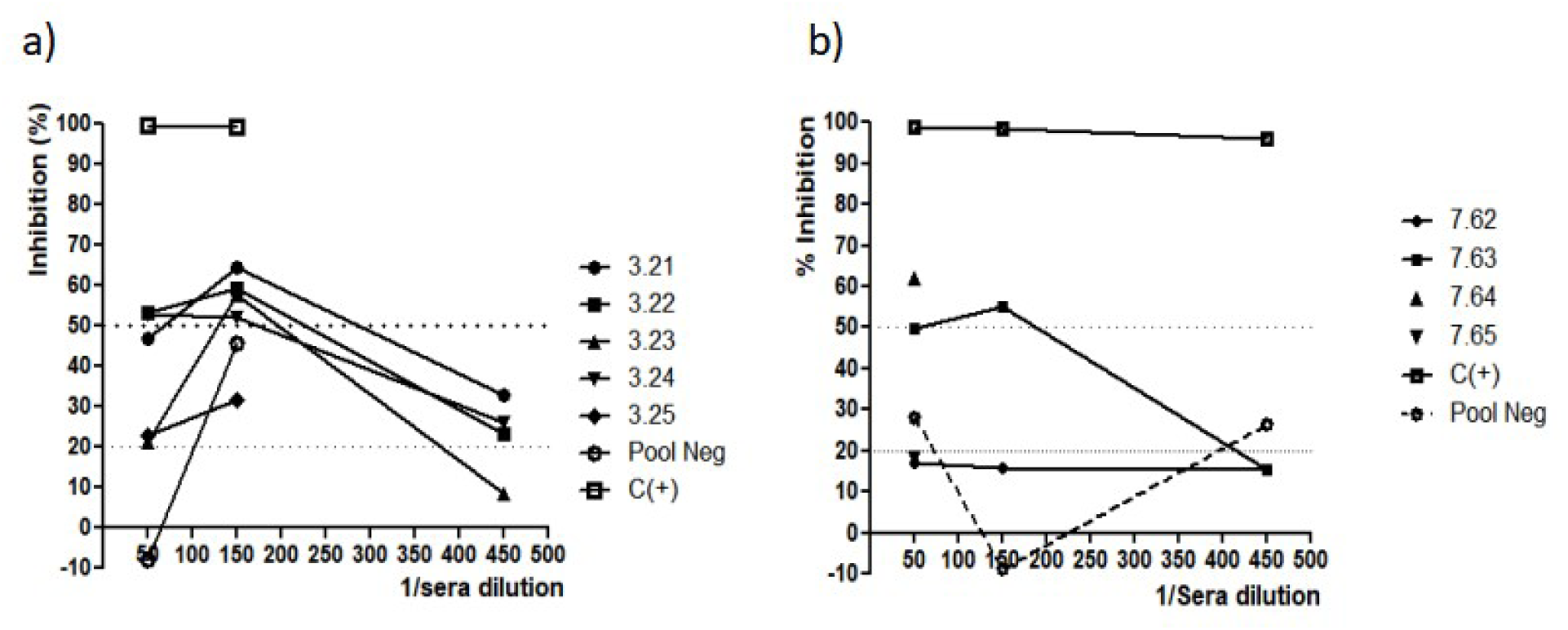
Neutralizing capacity of the antibody response generated in sera, measured by Ancestral strain Spike pseudotyped VSV assay. C57BL/6J mice were immunized with three doses administered at days 0, 7 and 21. The sera were collected 18 days after the last dose. a) mice immunized with bivalent formulation (N+ODN-39M+S2) by intranasal route, group 3 in the assay b) mice immunized with bivalent formulation (N+ODN-39M+S2+Alum) by subcutaneous route; group 7 in the assay. Pool Neg: pool of sera from Placebo immunized group, C(+): Pool of sera with known neutralizing titer.

#### 3.1.4 IFNγsecretion response generated in spleen after immunization with the bivalent formulation

The secretion of IFNγ by splenocytes of immunized mice was evaluated 26 days after the last dose. A positive response against S2 antigen was only observed for the group that received the bivalent formulation by intranasal route (Figure 4a). In this group, 4 out of 5 animals show low, but positive, levels of IFNγ secretion. On the other hand, the anti-N specific response was detected only for the groups intranasally immunized with the bivalent formulation or the control group receiving N+ODN-39M (Figure 4b). The response in this last group shows a trend to be higher.

**Figure 4.**
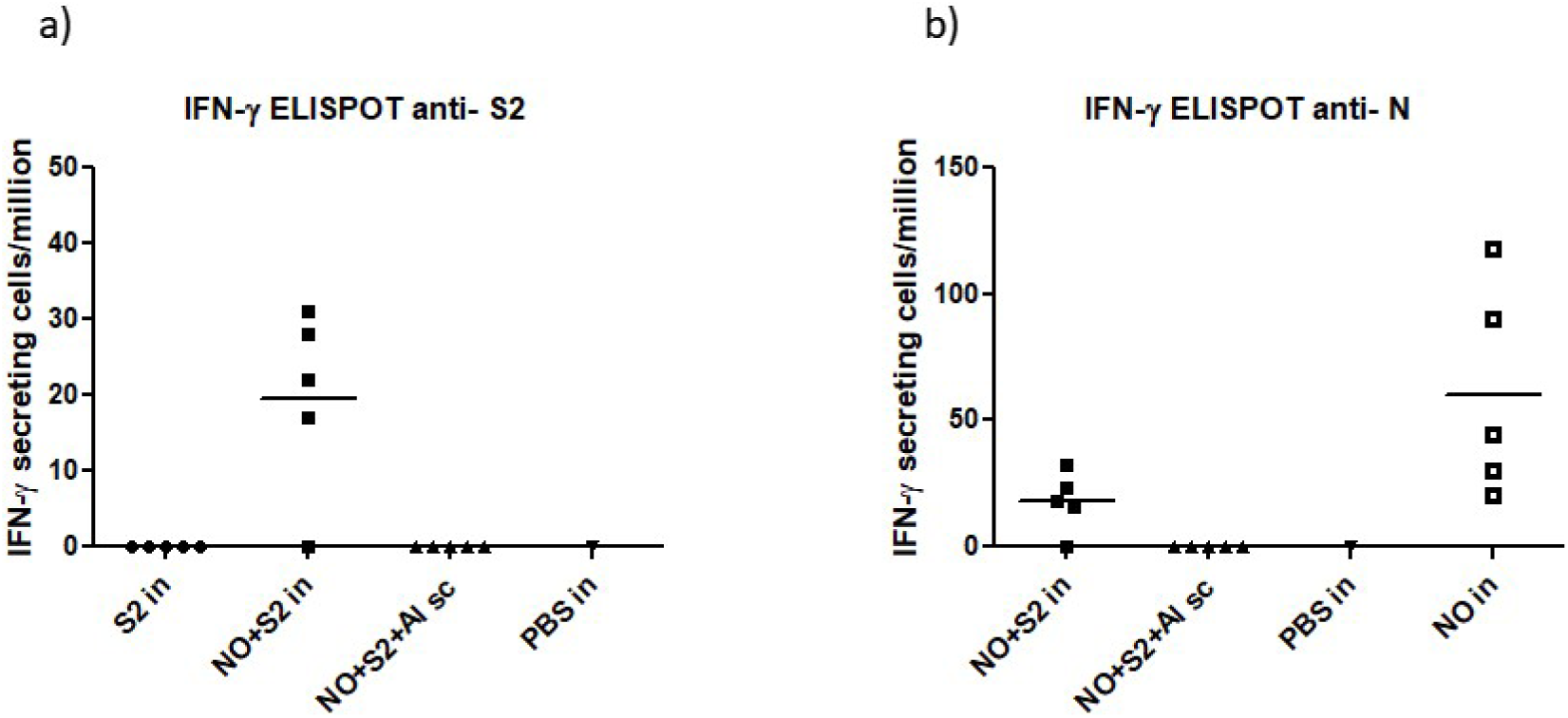
IFNγ secretion response measured, by ELISPOT, in spleens collected 26 days after the third dose. Anti-S2 (a) and Anti-N (b). C57BL/6J mice were immunized with three doses administered at days 0, 7 and 21. A dose of 10μg of S2 protein and 5μg of N protein per mouse was employed. Spleen samples from five individual mice per group were processed and stimulated with the SARS-CoV-2 antigens, S2 from Ancestral strain and N from Delta strain after 48h the IFNγ secretion was assessed.

### 3.2 Immunogenicity in Balb/C mice of the bivalent formulation N+ODN-39M+S2 intranasally administered

#### 3.2.1 Antibody response induced by the bivalent formulation (N+ODN-39M+S2)

Considering the results previously obtained in C57BL/6J mice and the relevance of develop a nasal vaccine candidate to fight respiratory infections we selected this administration route to further evaluations in Balb/C mice. Figure 5 shows the antibody response, IgG and IgA induced in sera and BAL, respectively, by the bivalent formulation.

**Figure 5.**
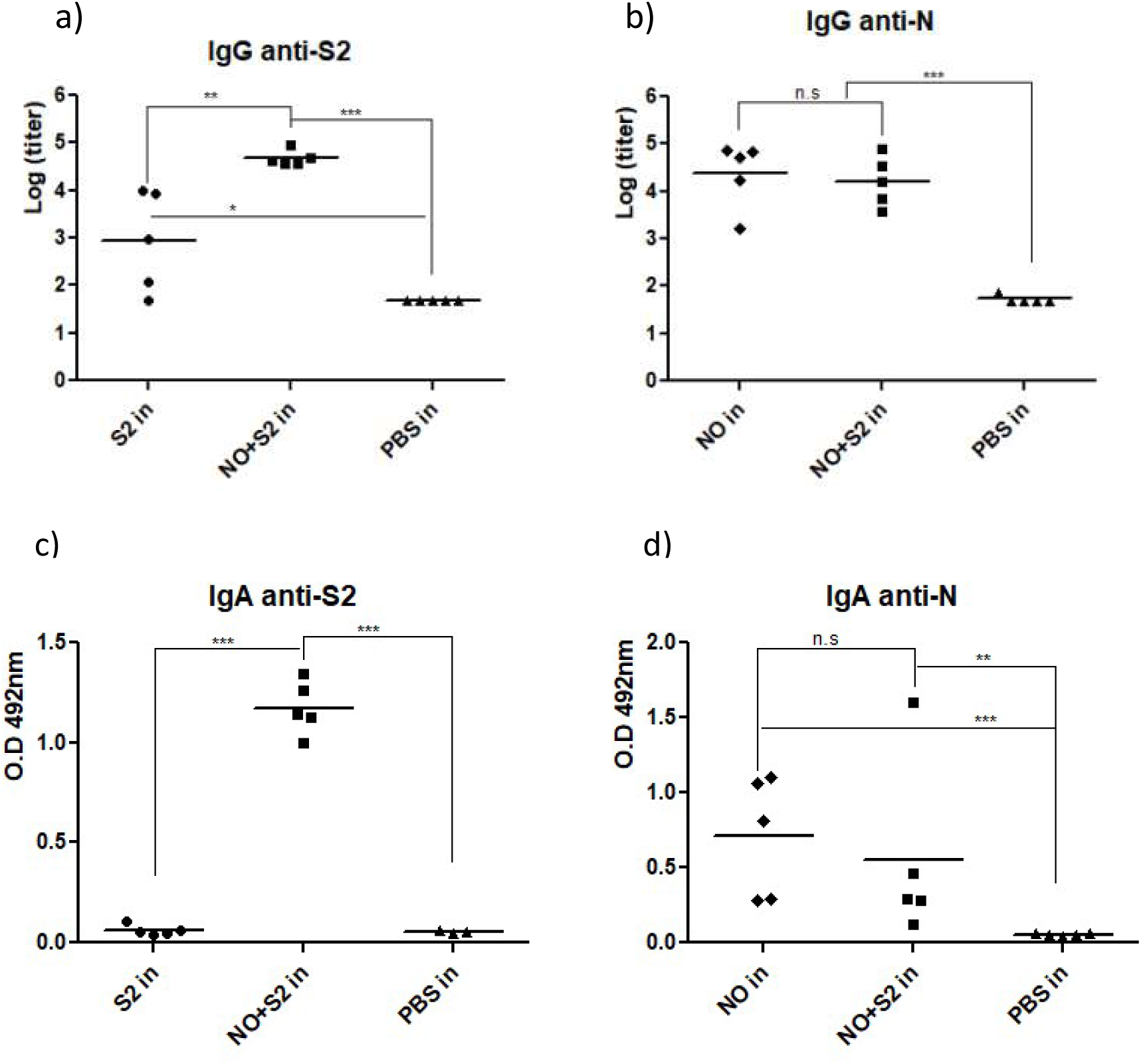
Antibody response induce by the bivalent formulation (N+ODN-39M+S2) administered by intranasal route. Balb/C mice were immunized with three doses administered at days 0, 15 and 30. A dose of 10μg of S2 protein and 5μg of N protein per mouse was employed. Thirty days after the last dose sera and BAL were collected and the IgG and IgA antibody response, respectively, was evaluated by ELISA. Anti-S2 IgG (a), IgA (c). Anti-N IgG (b), IgA (d). IgG was represented as arbitrary titers and IgA as O.D at 492nm.

The bivalent formulation was able to generate a high response of IgG anti-S2, which was significantly superior to the one generated by the administration of S2 protein in PBS (Figure 5a). Consistently with data from C57BL/6J studies, the S2 protein without adjuvants is able to induce IgG titers after nasal administration. However, only the bivalent formulation generated a detectable anti-S2 IgA response in BAL (Figure 5c). The pattern of IgG subclasses was also studied (Supplementary Figure 1). The bivalent formulation demonstrated the induction of a significantly higher response of both, IgG1 (p<0.01) and IgG2a (p<0.001) compared with the group receiving S2 in PBS. The titers of IgG2a anti-S2 generated were notably high. Furthermore, the neutralizing capacity was evaluated in sera for the bivalent formulation (Supplementary Figure 2). Consistently with the observed in the previous experiments for this formulation, the neutralizing response detected was low, almost completely negative, only the serum of 1 out 5 Balb/C mice showed the capacity to inhibit more than 50% at the lowest dilution assayed (1/50).

On the other hand, the IgG anti-N response induced in sera by the bivalent formulation was also high but similar to the induced by the control group that received N+ODN-39M (Figure 5b). Both groups showed a positive, and similar in magnitude, anti-N IgA response in BAL samples (Figure 5d).

#### 3.2.2 IFNγresponse induced by the bivalent formulation (N+ODN-39M+S2)

The bivalent formulation generated a positive IFNγ secretion response against N protein in 3 out 5 mice. However, none of the immunized animals showed a positive response against S2 protein, and only one mouse responded after stimulation with the N_351-365_ peptide (Figure 6). The control group receiving N+ODN-39M also showed a positive anti-N response, consistently with previous data, in this case all the evaluated animals responded. On the other hand, no response was detected for the group receiving S2 in PBS, neither the Placebo group receiving only PBS.

**Figure 6.**
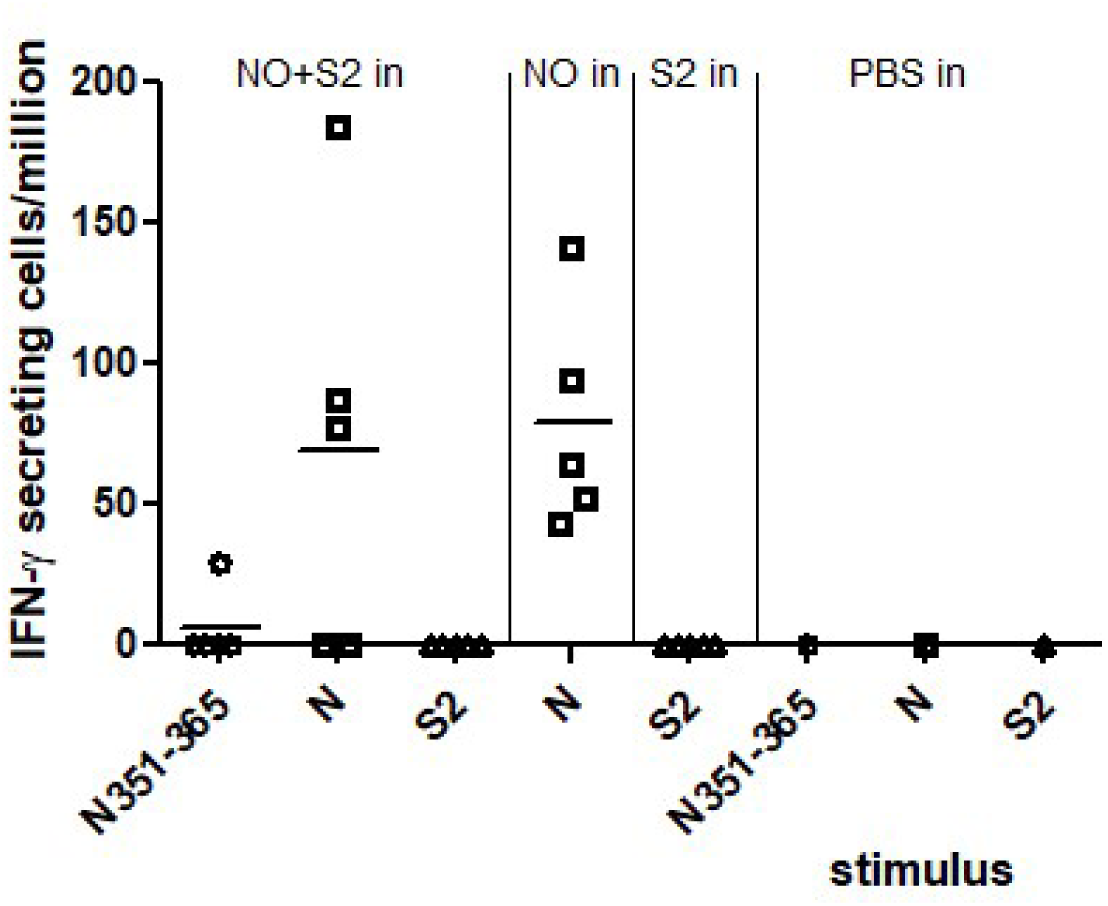
IFNγ secretion response measured, by ELISPOT, in spleens collected 30 days after the third dose. Balb/C mice were immunized with three doses administered at days 0, 15 and 30. A dose of 10 μ g of S2 protein and 5 μ g of N protein per mouse was employed. Spleen samples from five individual mice per group were processed and stimulated with the SARS-CoV-2 antigens, S2 from Ancestral strain and N from Delta strain, and the N_351-365_ conserved T CD4 peptide by 48h. NO: N+ODN-39M, in: intranasal.

### 3.3 Cross-reactive antibody response induced by the bivalent formulation (N+ODN-39M+S2) in two different mice strains

In Balb/C mice intranasally immunized with the bivalent formulation we found a positive recognition in sera for S2 proteins from SARS-CoV-1 and MERS-CoV, with a trend to a higher recognition observed against the SARS-CoV-1 protein (Figure 7a). The same sera samples are able to recognize, showing high and similar O.D values, the N proteins from SARS-CoV-2 Delta and Omicron strains, and from SARS-CoV-1 (Figure 7b). On the other hand, the cross-reactivity to S2 from SARS-CoV-1 and MERS-CoV was also observed in the sera of C57BL/6J mice intranasally immunized with the bivalent formulation (Figure 7c). In this mice strain, high and similar (p≥0.05) antibody titers were observed against the three S2 proteins evaluated (including S2 from SARS-CoV-2 Ancestral strain, SARS-CoV-1 and MERS-CoV), although a trend to a lower recognition was detected for S2 from MERS-CoV.

**Figure 7.**
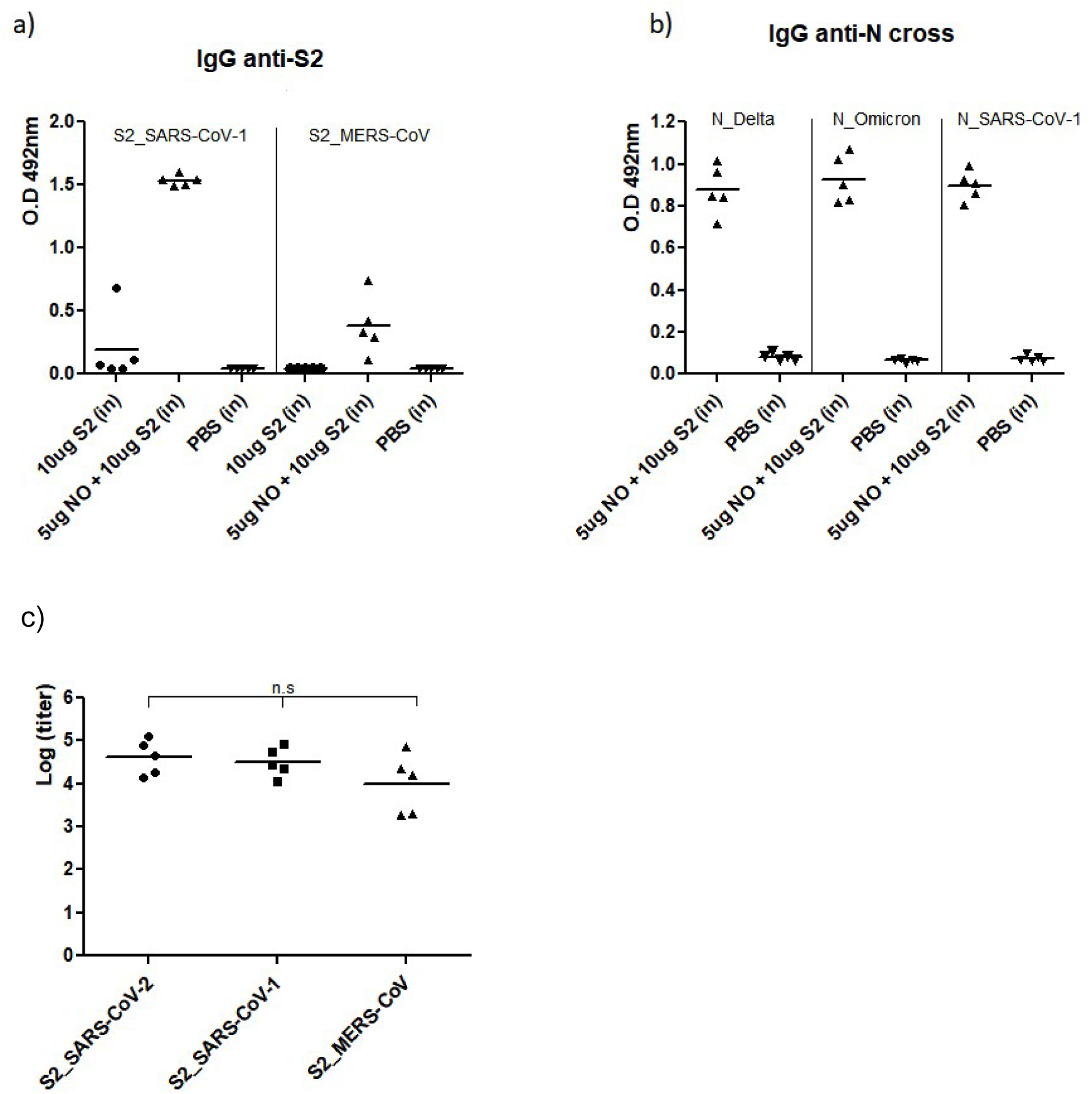
IgG cross-reactive response induced in sera by N+ODN-39M+S2 bivalent formulation. Balb/C and C57BL/6J mice were immunized with three doses administered at days 0, 15, 30, and 0, 7, 21, respectively. The recognition of proteins from different SARS-CoV-2 strains (S2 from Ancestral variant and N from Delta and Omicron variants), SARS-CoV-1 and MERS-CoV was evaluated by ELISA. a) anti-S2 cross-reactivity in Balb/C mice, sera samples diluted 1:100; b) anti-N cross-reactivity in Balb/C mice, sera samples diluted 1:1000; c) anti-S2 cross-reactivity in C57BL/6J mice.

The cross-reactive profile against both antigens, S2 and N, induced by the bivalent formulation was also evaluated in BAL samples coming from Balb/C and C57BL/6J mice intranasally immunized (Figure 8). Both mice strains showed a notable recognition of the S2 protein from SARS-CoV-1 by IgA antibodies in BAL samples (Figure 8a), a trend to develop a higher response was observed for Balb/C. At this mucosal samples the level of cross-reactivity against S2 from MERS-CoV was almost completely negative for both mice strains.

**Figure 8.**
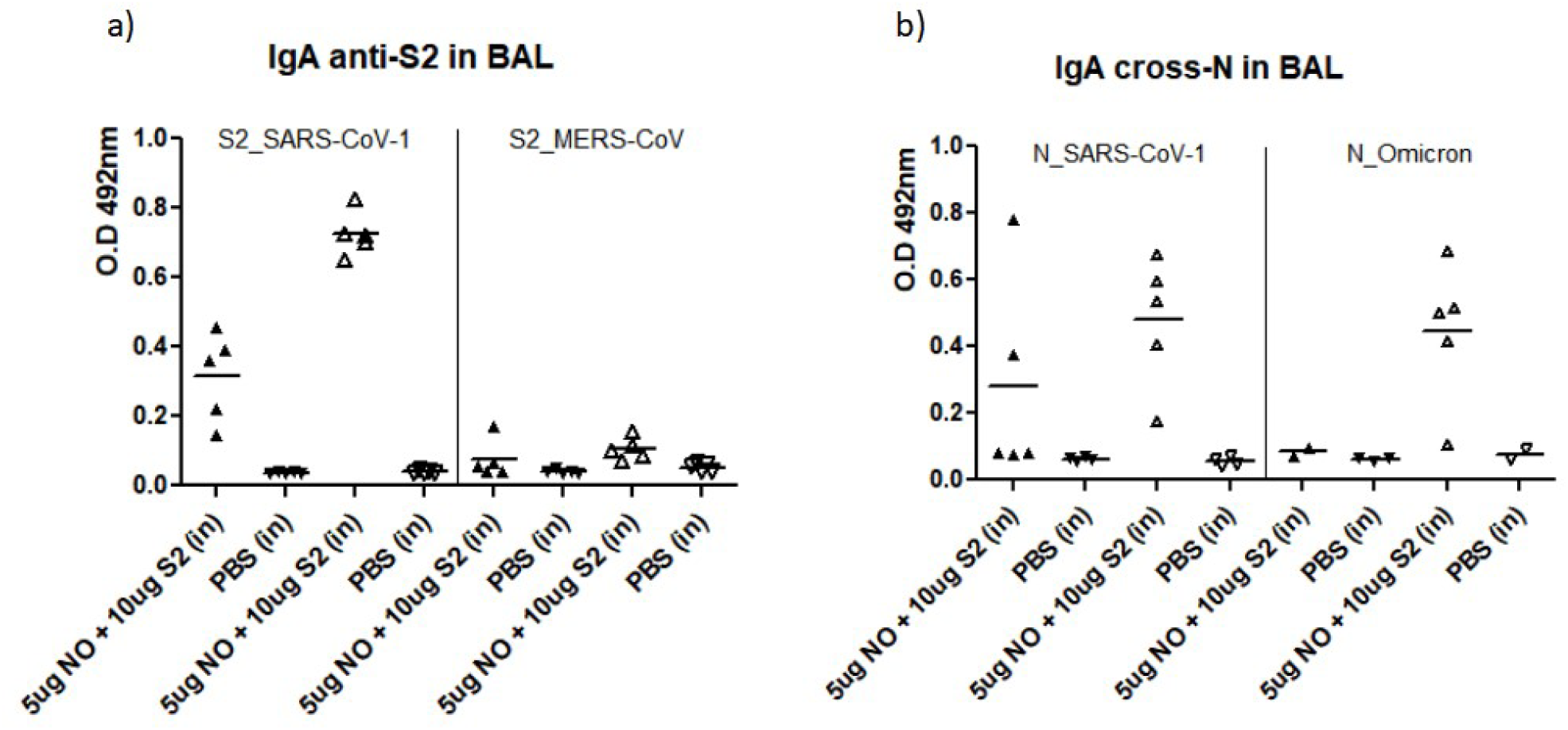
IgA cross-reactive response induced in BAL by the N+ODN-39M+S2 bivalent formulation. Balb/C (empty triangles) and C57BL/6J (filled triangles) mice were immunized with three doses administered at days 0, 15, 30, and 0, 7, 21 days, respectively. The recognition of S2 and N proteins from other SARS-CoV-2 strains, SARS-CoV-1 and MERS-CoV was evaluated by ELISA. BAL samples were evaluated without dilution. a) anti-S2 cross-reactivity; b) anti-N cross-reactivity.

Concerning the cross-reactivity against N protein, both strains of mice recognized the N protein from SARS-CoV-1, again a trend to develop a higher cross-reaction was detected for Balb/C mice (Figure 8b). In line, only immunized Balb/C animals showed a positive recognition for the N protein from SARS-CoV-2 Omicron strain. However, for this last assay only 2 sera samples were available in the C57BL/6J immunization group.

## 4. Discussion

The development of a second generation of vaccines with the ability to induce a broader scope of protection against SARS-CoV-2 variants and other potential emerging Coronavirus continues in the spotlight nowadays. Worldwide, there are several research groups, coalitions and initiatives focused in investigations related with pandemic preparedness strategies. Our group have been working in the field and published several papers related with the development of vaccine candidates for nasal administration based on the selection of Coronavirus conserved proteins or regions as key point to induce a broad scope of cross-reactivity and protection [**21, 22, 24**]. One of the previous vaccine candidates under development was based on a chimeric protein that includes two selected regions from SARS-CoV-2, one belonging to the Spike S2 subunit (amino acids 800-1020 in post-fusion fiber structure) and the other corresponding with the C-terminal domain of the nucleocapsid protein (amino acids 248-371) [**24**]. Considering that this candidate was unable to induce in mice the expected immune response against the S2 fragment, and also taking into account the relevance of S2 protein as target for more universal vaccines versus Coronavirus [**11-15**], we decided to explore in the present work a vaccine preparation containing the full length S2 (amino acids 708-1207, stabilized in pre-fusion conformation) and N proteins administered by intranasal route. These two protein antigens were selected by their high conservation between Sarbecovirus and other Coronavirus [**25, 26**], which will impact in the generation of a broad scope immune response. S2 protein also has an important role during the infection process and is target of neutralizing antibodies [**26, 27**]. However, the use of S2 recombinant protein as vaccine antigen choice has been hampered by its conformation instability [**28**]. On the other hand, the vaccine preparation under evaluation also includes the ODN-39M as CpG ODN adjuvant. This adjuvant has been previously evaluated as component of our nasal vaccine candidates showing a potent immuno-enhancing capacity and imprinting a bias to develop a Th1 pattern of immune response [**22, 24**].

The bivalent formulation, containing N+ODN-39M+S2, was evaluated in two different mice strains with the aim to expand the characterization of the induced immune response and cross-reactive profile. In C57BL/6J mice we explored two different administration routes, intranasal and subcutaneous. For the last route, Alum was incorporated as adjuvant in the preparation with the aim to generate a potent immune response and to use this group as comparative for the intranasal administered variant. The results indicate that a similar IgG antibody response, against S2 and N proteins, is generated in sera after the administration of the bivalent formulation by both administration routes. However, the intranasal administration favors the induction of a higher IgG2a response versus both proteins. In mice models, the production of IgG2a subclass is associated with the development of a Th1-like pattern of immune response [**29**]. Remarkably, the bivalent formulation administered by intranasal route, differing from the administered subcutaneously, shows the capacity to induce an IgA response in BAL, and an IFNγ secretion response by splenocytes, against both proteins included in the preparation. The generation of antibody response at mucosal sites, in this case the lower respiratory tract, represents a critical advantage to face respiratory infections [**30**]. The absent, or very low levels, of this kind of immune response induced by the classical COVID-19 vaccines constitutes a weakness with implications in the subsequent infections and the continue SARS-CoV-2 transmission and circulation [**30, 31**].

Specifically, the induction of anti-S2 antibody response in both, systemic and mucosal compartments is a very important result. It is known that some anti-S2 antibodies has viral neutralization capacity [**13-15**] and also other relevant effector functions, like ADCC, has been described and associated with protection [**11, 32**]. In this line, an interesting and unexpectedly finding of our experiments was the immunogenicity found for the S2 protein, without adjuvants, when is intranasal administered to C57BL/6J mice. A previous report [**33**] shows that in mice the SARS-CoV-2 Spike protein intranasally administered without adjuvants was unable to induce significant levels of IgG at systemic compartment, neither antibody or T cell mediated response at airway mucosal tissue. This data correlates with our own former results evaluating the SARS-CoV-2 RBD from Delta strain without adjuvants by intranasal route [**22, 24**]. In general, to induce an immune response by intranasal route it is required to use specific adjuvants (ex/ toxin derivatives, CpG ODN) or more complex delivery systems (ISCOMs, liposomes, nanovesicles) that allow to overcome the mucosal tissue natural barriers and favor the interaction with the antigen presenting cells [**34, 35**]. We hypothesized that in the case of S2 protein, its specific nature (including primary sequence and conformational flexibility) could favor its interaction with the antigen presenting cells from the nasal mucosa. The employed recombinant S2 protein, exposed at physiological nasal mucosa environment (pH, presence of proteases), could be, somehow, mimicking the natural S2 protein behavior, characterized by conformational changes and binding motifs (including the fusion peptide, heptad repeats (HR1 and HR2), transmembrane domain, and cleavage sites) that work together to mediate membrane fusion between the virus and the host cell [**14**].

On the other hand, the cell-mediated immune response is also regarded as relevant in Coronavirus protection. Specifically, in the case of SARS-CoV-2 several studies highlight the contribution of nucleocapsid-specific T cells in the protection [**36, 37**]. In this line, Matchett et al., 2021, demonstrated that the N protein, presented in adenovirus vectors, induced humoral and cellular immune response in mice, correlating with protection to SARS-CoV-2 challenges [**37**]. In another approach, Dangi et al., 2021, combines the spike and nucleocapsid proteins in a vaccine and observed the improvement of distal protection in the brain [**38, 39**]. Furthermore, N protein is also a target of cross-reactive memory T cells, which were associated with protection from infection in SARS-CoV-2 naïve contacts, thereby supporting the inclusion of N protein antigen in new vaccines generation [**36**]. On the other hand, the role of nucleocapsid-specific humoral immunity has been studied [**40**]. Dangi et al, 2022 [**39**] reported that mice receiving anti-N antibodies show an enhanced control of SARS-CoV-2. They demonstrated that nucleocapsid-specific sera are able to elicit antibody-dependent cellular cytotoxicity (ADCC) against infected cells. In humans, anti-N sera have been used to treat COVID-19 patients with promising results reducing the severity of the disease [**41**]. In addition, another recent work indicates that nucleocapsid serostatus prior to SARS-CoV-2 breakthrough infection correlates with disease protection by showing that anti-N seropositive individuals have an increased rate of virus clearance and lower peak viral loads compared to seronegatives [**42**].

Considering the results obtained in C57BL/6J mice strain, we carried out further evaluations of the nasally administered bivalent formulation in Balb/C mice. In this second mice strain the bivalent formulation consistently induced a high IgG antibody response against both SARS-CoV-2 proteins, S2 and N. In a similar way the presence of IgA response in BAL samples was detected against both proteins. Although the antibody titers induced against S2 protein were high in both mice strains, when the neutralizing capacity in sera was measured, using a Spike-pseudotyped VSV assay, the results were low or negative. A similar scarce neutralizing capacity was obtained in the sera from the C57BL/6J mice immunized subcutaneously with the bivalent formulation plus Alum. This last data suggests that the inability of the bivalent formulation to induce neutralizing antibody response is not related with the immunization route employed. Previously other authors has also failed in the induction of neutralizing antibodies using S2 protein as vaccine antigen [**13, 14**]. In our case, one plausible explanation could be related with the folding of the S2 protein used as immunogen, which is stabilized in the pre-fusion conformation. It is known that the conformational status of the S2 subunit significantly impacts its immunogenicity [**28**], with the post-fusion state being more effective at eliciting neutralizing antibodies. This behavior is based on the capacity of the post-fusion conformation to present critical conserved epitopes that are otherwise inaccessible in the pre-fusion state [**43**]. However, to reach by engineering an stable S2 trimeric protein conformation where the neutralizing epitopes are exposed continues to be a hot-spot [**28, 43**]. The impact of the S2 protein conformation on the induction of neutralizing antibody response is supported by the work of Pang et al [**14**]. They found the induction of neutralizing antibodies in rabbit and rhesus macaques immunized with a vaccine candidate based on HR1 and HR2 fusion protein that mimics the fusion intermediate conformation. However, other authors has also observed the induction of neutralizing antibodies in mice using prefusion stabilized S2 antigens [**44**].

On the other hand, it has been extensively demonstrated that the S1 subunit, specifically the RBD region, constitutes the major target for neutralizing response in SARS-CoV-2, while S2 protein represent a minority target of this kind of response [**2, 5**]. In addition, the differences between the animal models, vaccine preparations, and immunization routes, employed could explain the mismatch of our data with other published results [**14, 44**]. Another point that could explain the low neutralizing response observed in our work is the use of a pseudoviral system assay instead of real SARS-CoV-2 virus. The relevance and correlation of the pseudoviral based assays to measure the neutralizing capacity has been established before [**45, 46**], but mainly using sera generated against the full length Spike or RBD proteins. Considering the current neutralization results, and the other potential kind of effector functions that anti-S2 immunity can exert, the real effect in protection of the immune response generated after intranasal administration of the bivalent formulation N+ODN-39M+S2 should be evaluated using *in vivo* challenge experiments. The pending realization of this kind of studies constitute a limitation of the present work.

On the other hand, regarding the IFNγ secretion response measured in spleen cells, in Balb/C mice a positive response was only detected after stimulation with N protein (3 out 5 mice) and N_351-365_ conserved peptide (1 out 5 mice). No S2-specific IFNγ secretion was observed differing from the previous data coming from C57BL/6J mice. This discrepancy could be explained by immune system differences between both mice strains. It is known that C57BL/6 mice strain is more prone to develop a Th1-like cellular immune response while Balb/C is considered as more Th2-biased [**47**]. Furthermore, considering the role of mucosal resident T cell response in the protection against viral infections [**48**], future studies should explore the potential induction of cell-mediated immune response at mucosal compartment. Recently, our group have obtained clear evidences of the generation of this type of response in lungs from mice immunized with other similar nasal vaccine preparation containing N+ODN-39M (unpublished data).

Altogether, in both mice strains a clear adjuvant effect exerted by the mix N+ODN-39M over the S2 protein was demonstrated by intranasal route. This adjuvant capacity, with a bias to a Th1 response, had been previously demonstrated by our group in a nasal formulation containing RBD protein as inducer of neutralizing antibodies instead of S2 protein [**22**]. In our opinion, the nanometric particle structure demonstrated for the recombinant N protein from SARS-CoV-2, which is reinforced after the mix with the ODN-39M [**21, 22**] could explain the high immunogenicity of the bivalent formulation by intranasal route. Particulate preparations, virus-like or capsid-like, with a size in the nanometric range (preferentially between 20-80 nm) are able to interact efficiently with the mucosal antigen presenting cells to develop a relevant immune response [**49, 50**]. In addition, the ODN-39M, a CpG ODN with a complete phosphodiester backbone is able to activate TLR9, with is highly represented in plasmacytoid dendritic from nasal mucosa [**51**].

Finally, we studied the scope of cross-reactivity generated in sera and BAL from C57BL/6J and Balb/C mice intranasally immunized with the bivalent formulation. In both mice strains we found that the IgG antibodies induced at systemic level were able to recognized the S2 proteins from SARS-CoV-1 and MERS-CoV, and the N protein from different SARS-CoV-2 strains and from SARS-CoV-1. However, at mucosal compartment the cross-reactivity of the IgA antibody response detected in BAL covers until SARS-CoV-1 for both proteins, showing only very slightly evidences of recognition for the S2 from MERS-CoV. These results, in general, are in line with previous reports about the capacity of S2 protein to induce cross-reactivity and even cross protection against other Betacoronavirus [**11, 12, 32**]. In the case of the results of anti-N cross-reactivity the data found in this study match with other previous studies published by our group [**21, 22**], and is in line with the percentage of homology between N proteins from different Coronavirus. Within the Sarbecovirus subgenus, it is estimated a range of 87–99% of identity whereas, for N MERS-CoV, the identity with the nucleocapsid of SARS-CoV-2 is 48% [**25**].

In conclusion, to our knowledge, this is the first study exploring the immunogenicity by intranasal route of the full-length prefusion S2-only protein, without any carrier system or delivery platform (viral vector or mRNA). The immune response induced by the nasal bivalent formulation containing the full-length N and S2 proteins from SARS-CoV-2 results promising, pointed out this vaccine candidate as another appealing formulation with a potential broader scope of protection against Betacoronavirus. The generation of cross-reactive antibody response also at lower respiratory tract level constitutes a very relevant feature, suggesting the potential use of this candidate as booster option in current population. Future protection studies must be done to demonstrate the real effect of the generated immunity.

## Author Contributions

Conceptualization, L.H., Y.L. (Yadira Lobaina), E.S., G.G., and Y.P.; Supervision, L.H., Y.P., G.G. and W.L.; Investigation, Y.L. (Yadira Lobaina), R.C., D.V-B., M.Z., Z.Z., Y.L. (Yaqin Lan); Formal analysis, L.H., Y.L. (Yadira Lobaina) and Y.P.; Funding acquisition, L.H. and Y.P.; Writing—Original Draft, Y.L. (Yadira Lobaina) and L.H.; Writing—Review & Editing, Y.L. (Yadira Lobaina), D.V-B.; Project administration, L.H., W.L. and Y.P. All authors have read and agreed to the published version of the manuscript.

## Funding

This work was supported by MOST “National key R&D program of China (2021YFE0192200)”, “PNCT CITMA, Cuba”, “Hunan Provincial Base for Scientific and Technological Innovation Cooperation (2019CB1012)”, “The Science and Technology Innovation Program of Hunan Province, (2020RC5035)”, “Hunan Provincial Innovative Construction Program (2020WK2031).

## Institutional Review Board Statement

The animal study protocols were approved by the Institutional Animal Care and Use Committee at Beijing Vital River Laboratory Animal Technology Co., Ltd. and Hunan PRIMA Animal facility. The standard of laboratory animal room complied with the national standard of the people’s Republic of China GB14925-2010. Protocols: PANCOV03 (approved on March 2022) and HNSE2023(3)049 (approved on July 2023).

## Supplementary Figures

**Supplementary Figure 1.**
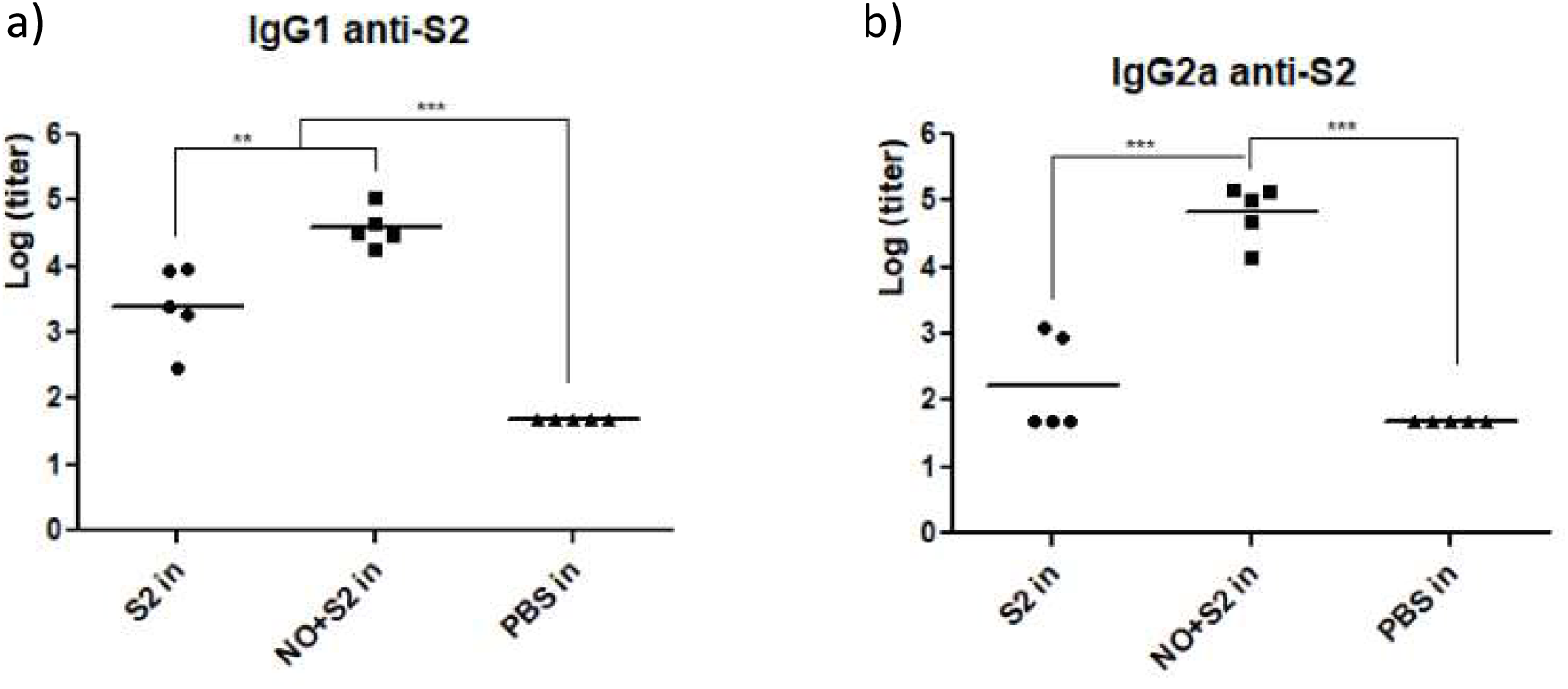
Anti-S2 IgG subclasses in sera after N+ODN-39M+S2 intranasal administration in Balb/C mice. Mice were immunized with three doses administered at days 0, 15 and 30. A dose of 10μg of S2 protein and 5μg of N protein per mouse was employed. Thirty days after the last dose sera were collected and the IgG subclasses response was evaluated by ELISA. a) IgG1, b) IgG2a.

**Supplementary Figure 2.**
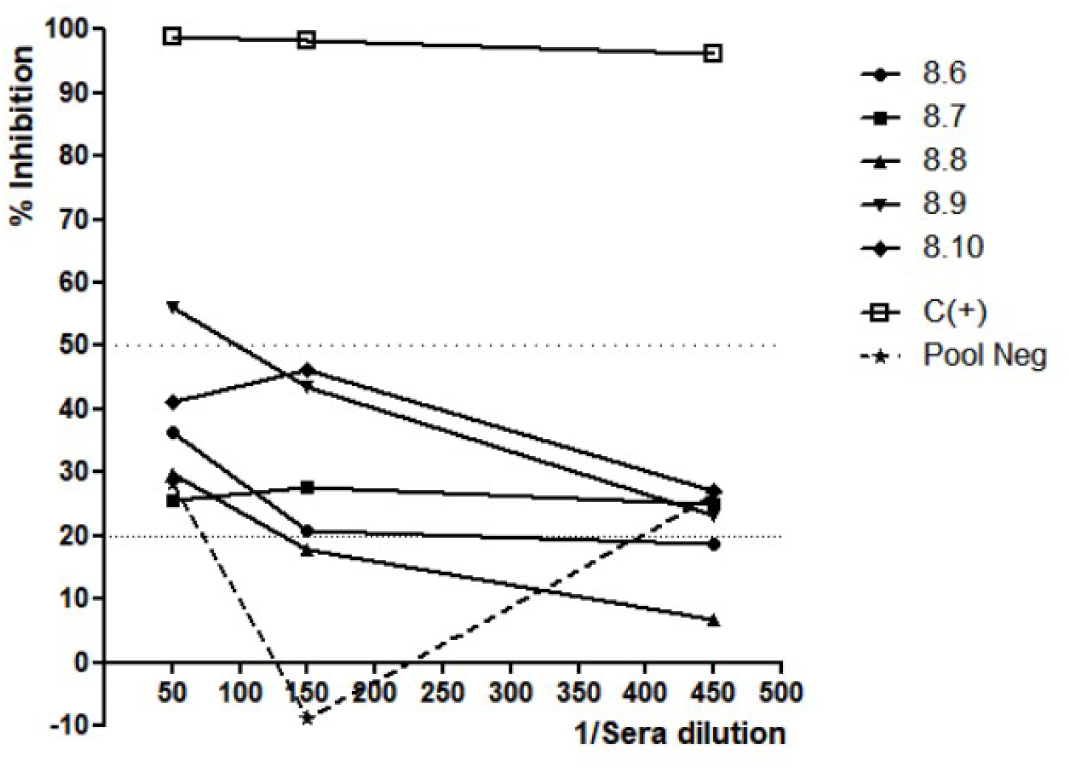
Neutralizing capacity in sera after N+ODN-39M+S2 intranasal administration in Balb/C mice. Mice were immunized with three doses administered at days 0, 15 and 30. A dose of 10μg of S2 protein and 5μg of N protein per mouse was employed. Thirty days after the last dose sera were collected and the neutralizing capacity measured using Ancestral strain Spike pseudotyped VSV assay.

## Notes

### Competing Interest Statement

The authors have declared no competing interest.

